# Response of soil microbial communities to mixed forests of European beech and conifers: Variations with site conditions

**DOI:** 10.1101/2020.07.21.213900

**Authors:** Jing-Zhong Lu, Stefan Scheu

**Affiliations:** J. F. Blumenbach Institute of Zoology and Anthropology, University of Göttingen, Untere Karspüle 2, 37073 Göttingen, Germany; Center of Biodiversity and Sustainable Land Use, University of Göttingen, Büsgenweg 1, 37077 Göttingen, Germany

**Keywords:** Douglas-fir, Norway spruce, soil respiration, microbial stress, ecosystem functioning, PLFA

## Abstract

Tree - soil interactions depend on environmental conditions. Planting trees may strongly impact microorganisms in particular at unfavorable site conditions, compromising the functioning of soil microorganisms. To understand the effects of tree species composition on soil microorganisms, we quantified structural and functional responses of soil microorganisms to forest types across environmental conditions using substrate-induced respiration and phospholipid fatty acid analyses. Five forest types were studied including pure stands of native European beech (*Fagus sylvatica*), range expanding Norway spruce (*Picea abies*), and non-native Douglas-fir (*Pseudotsuga menziesii*), as well as the two conifer - beech mixtures. We found that microbial functioning strongly depends on site conditions, in particular on soil nutrients. At nutrient-poor sites, soil microorganisms were more stressed in pure and mixed coniferous forests, especially in Douglas-fir, compared to beech forests. By contrast, microbial structure and functional indicators in beech forests varied little with site conditions, likely because beech provided high amounts of root-derived resources for microbial growth. Since soil microbial communities are sensitive to Douglas-fir, planting Douglas-fir may compromise ecosystem functioning in particular at nutrient-poor sites. Overall, root-derived resources are important for determining the structure and functioning of soil microbial communities, and soil microorganisms sensitively respond to plantations containing tree species that may differ in the provisioning of these resources.

## 1. Introduction

Trees affect soil microorganisms through several pathways, most importantly by litter and root exudates (Wardle et al., 2004; Högberg and Read, 2006). These resources are likely to shape microbial community composition because dissimilar carbon resources favor different guilds of microorganisms (Lindahl et al., 2007; Fanin et al., 2019). Despite that microbial community composition may change in response to variations in tree species composition, the relative role of litter and root exudates in structuring soil microbial communities remains controversial (Bluhm et al., 2019). The typically thicker organic layers in coniferous compared to deciduous forests have been attributed to the recalcitrance of needles (Augusto et al., 2015), but root and mycorrhizal fungi also affect soil carbon storage (Clemmensen et al., 2013; Averill et al., 2014). Studying the linkage between trees and soil microorganisms may allow uncovering the pathways by which trees drive microbial community composition.

Planting coniferous trees beyond their native range has become common worldwide (Castro-Díez et al., 2019). Although providing timber and other benefits to humans, conifers may detrimentally affect soil microorganisms, compromising carbon and nutrient cycling (Berger and Berger, 2012; Castro-Díez et al., 2019). To improve microbial functioning, admixing conifers to deciduous forests has been suggested as it may increase resource availability to soil microorganisms in mixed litter (Hättenschwiler et al., 2005; Cremer et al., 2016). Further, broadleaf mixed forests are likely to increase both aboveground and belowground biomass due to interspecific facilitation and improved resource partitioning, resulting in higher resource availability to microorganisms (Cardinale et al., 2007; Emmett Duffy et al., 2017). The importance of mixed stands for ecosystem functioning is increasingly recognized, particularly under unfavorable environmental conditions (Ratcliffe et al., 2017; Wright et al., 2020). However, it is little understood how effects of tree species composition on soil microorganisms vary with site conditions, such as soil nutrient status (Malchair and Carnol, 2009).

Facilitation among plants is more pronounced in nutrient-poor soil according to the stress-gradient hypothesis, and such positive interspecific interactions are likely to be mediated by soil microorganisms (Defossez et al., 2011; David et al., 2020). Plants do not passively tolerate environmental stress, but respond in various ways to unfavorable growth conditions, such as by shifting the allocation of resources towards roots in nutrient-poor soil (Callaway et al., 2003; Yan et al., 2016). Recently, de Vries et al. (2019) has shown that under stress conditions herbaceous species change the quality of root exudates and induce higher microbial respiration, presumably facilitating nutrient capture by stimulating microorganisms. Phenotypic plasticity or morphological changes induced by the environment may differ between species, exacerbating differences in resource availability to soil microorganisms under nutrient-poor conditions (Meier and Leuschner, 2008; Schall et al., 2012). Thus, the impact of tree species composition on microbial community composition and functioning may be more pronounced at nutrient-poor than nutrient-rich sites.

A better understanding of the effects of tree species composition on soil microorganisms has implications for forest management, especially in temperate and boreal regions where tree species richness is low and managed forests are dominated by one or a few species (Knoke et al., 2008; Bauhus et al., 2010). European beech is the climax species in lowland and lower montane regions in Central Europe (Leuschner et al., 2017). The most popular timber species, Norway spruce originally occurred in higher mountain ranges and boreal regions, but has been planted widely in lowlands (Knoke et al., 2008). In recent years, Norway spruce has been damaged severely by extreme weather and bark beetle outbreaks, events predicted to become more frequent in the future (Pettit et al., 2020). Although the admixture of Norway spruce to native European beech forests may reduce the risk of damage while maintaining economic gains, Douglas-fir is increasingly planted (Schmid et al., 2014). Since its introduction from North America over 150 years ago, Douglas-fir has become the most abundant non-native cultivated tree species in Central Europe (Schmid et al., 2014). To date, the impact of planting Douglas-fir on biodiversity and functioning of forests is little studied, and this applies in particular to the belowground system, although e.g., Douglas-fir has been suggested to affect soil chemistry in a similar way than spruce (Prietzel and Bachmann, 2012; Schmid et al., 2014). Overall, we lack a comprehensive evaluation of forest types on soil microorganisms across soil nutrient conditions.

Here, we studied microbial community composition and functioning in litter and soil of five forest types of different soil-nutrient status. Forest types included pure stands of European beech, Norway spruce, Douglas-fir, and the two conifer - beech mixtures. We analyzed the structure of microbial communities using phospholipid fatty acid patterns, and their functioning using microbial basal respiration, biomass and stress indicators. In general, we assumed the structure of microbial communities to be closely linked to their functioning. In particular, we hypothesized that (1) coniferous trees will more detrimentally affect microbial structure and functioning compared to European beech, with the effects being similar in Norway spruce and Douglas-fir forests, and intermediate in mixed forests. Further, we hypothesized that (2) microbial community structure and functioning will be strongly impacted by forest type at nutrient-poor sites, but less at nutrient-rich sites. Since the litter layer is less buffered against environmental harshness than the organic and mineral soil, we also hypothesized that (3) forest types will affect soil microorganisms more strongly in litter than in soil.

## 2. Methods

### 2.1. Field sites

The study included five forest types arranged as quintets at eight sites in northern Germany, covering a range of environmental conditions (5 forest types × 8 sites; Fig. S1). The four sites in the south stock on fertile soil and receive higher precipitation. Mean annual precipitation is 821–1029 mm. Parental rock is either loess influenced Triassic sandstone or Paleozoic shares of greywacke, sandstone, quartzite and phyllite, resulting in soil types of partly podsolic Cambisol and Luvisol. The four sites in the north are located on out-washed sand with the soil type of Podzol. The mean annual precipitation is 672–746 mm. The southern sites are richer in nutrients than the northern sites as reflected by higher total soil P, cation exchange capacity and pH; more details on site characteristics and soil chemical properties are given in Table 1 (Ammer et al., 2020; Foltran et al., 2020). Hereafter, we refer to the southern sites as nutrient-rich sites and to the northern sites as nutrient-poor sites.

**Table 1.** General information as well as soil physical and chemical properties of the study sites. Soil texture is based on 5–30 cm mineral soil, and soil chemical properties are based on 0–5 cm mineral soil. Soil moisture is expressed as percentage of dry mass, and was measured in litter, 0–5 and 5–10 cm soil. Ca2+ concentration was analyzed in exchangeable form by NH4Cl extraction, and total P was determined by pressure digestion (for detailed see Methods and Foltran et al. 2020).

| Site condition |  | Rich sites |  |  |  | Poor sites |  |  |  |
| --- | --- | --- | --- | --- | --- | --- | --- | --- | --- |
| Site number |  | 1 | 2 | 3 | 4 | 5 | 6 | 7 | 8 |
| General information | Site name | Harz | Dassel | Winnefeld | Nienover | Nienburg | Unterlüß | Göhrde II | Göhrde I |
|  | Latitude (°N) | 51.770 | 51.726 | 51.662 | 51.696 | 52.621 | 52.837 | 53.127 | 53.201 |
|  | Longitude (°E) | 10.397 | 9.704 | 9.574 | 9.526 | 9.281 | 10.331 | 10.814 | 10.801 |
|  | MAT (°C) | 7.63 | 8.59 | 8.90 | 9.01 | 9.70 | 9.03 | 9.19 | 9.20 |
|  | MAP (mm) | 1029.24 | 821.29 | 822.43 | 874.97 | 733.34 | 746.56 | 681.68 | 672.63 |
|  | Slope (°) | 15.52 | 2.96 | 4.97 | 6.56 | 1.85 | 1.19 | 1.43 | 2.05 |
|  | Elevation (m) | 510.80 | 426.17 | 349.06 | 323.37 | 92.00 | 161.20 | 129.80 | 120.00 |
| Texture & moisture | Sand% | 16 | 26 | 20 | 20 | 80 | 79 | 79 | 73 |
|  | Silt% | 16 | 53 | 57 | 57 | 13 | 15 | 15 | 24 |
|  | Clay% | 68 | 21 | 23 | 23 | 7 | 6 | 6 | 3 |
|  | Water% (Litter) | 237.2 | 221.2 | 235.5 | 232.4 | 245.9 | 303.1 | 328.5 | 265.7 |
|  | Water% (0–5) | 135.5 | 106.6 | 84.7 | 62.7 | 71.4 | 120.2 | 88.1 | 76.2 |
|  | Water% (5–10) | 59.1 | 39.6 | 37.8 | 29.6 | 22.4 | 26.9 | 23.9 | 22.9 |
| Chemical properties | C % | 7.09 | 6.19 | 7.50 | 5.30 | 7.97 | 4.03 | 9.31 | 5.36 |
|  | N % | 0.37 | 0.34 | 0.37 | 0.29 | 0.28 | 0.17 | 0.34 | 0.24 |
|  | C/N | 19.16 | 18.21 | 20.27 | 18.28 | 28.46 | 23.71 | 27.38 | 22.33 |
|  | CEC (mmol dm<sup>-3</sup>) | 237.31 | 121.26 | 114.91 | 95.96 | 74.81 | 47.79 | 71.10 | 67.72 |
|  | pH (KCl) | 3.42 | 3.15 | 3.34 | 3.47 | 3.00 | 3.00 | 2.71 | 3.01 |
|  | P (mg kg<sup>-1</sup>) | 0.61 | 0.54 | 0.44 | 0.46 | 0.16 | 0.11 | 0.17 | 0.24 |
|  | Ca<sup>2+</sup> (CEC; mmol dm<sup>-3</sup>) | 28.35 | 13.77 | 32.56 | 27.88 | 20.38 | 12.91 | 7.90 | 11.28 |

MAT: mean annual temperature
MAP: mean annual precipitation
CEC: cation exchange capacity

Each site comprised pure stands of European beech (*Fagus sylvatica* L.; Be), Douglas-fir (*Pseudotsuga menziesii* [Mirbel] Franco.; Do) and Norway spruce (*Picea abies* [L.] Karst.; Sp), as well as two conifer-beech mixtures (Douglas-fir/European beech and Norway spruce/European beech; Do/Be and Sp/Be). On average, trees were more than 50 years old. Within sites, the distance between stands ranged from 76 m to 4600 m, and the distance between sites ranged from 5 to 190 km. Within each stand, plots of 2500 m^2^ were established, mostly in rectangular shape (50 × 50 m). Tree species in pure stands comprised more than 90% of the total basal area, and that in mixed stands 33–53% for beech and 53–60% for conifers.

### 2.2. Soil sampling and physio-chemical analyses

Samples were taken between November 2017 and January 2018. Soil cores of 5 cm diameter were taken by a metal cylinder, and were separated into litter, 0–5 and 5–10 cm soil depth. In each plot, three cores spaced by 5 m were taken. Samples from the same depth were pooled, resulting in 120 samples (40 plots × 3 depths). The soil was sieved through 2 mm mesh, and the litter was cut into pieces (< 25 mm^2^) (Maraun and Scheu, 1995). Roots > 2 mm in diameter and stones were removed. Samples were stored at −20°C. The pH was determined using a ratio of sample to solution (g/ml; KCl, 1 M) of 1:10 for litter, 1:5 and 1:2.5 for 0–5 and 5–10 cm soil, respectively. Moisture content was measured by drying samples at 105°C for 48 h. Water content did not differ consistently between nutrient-rich and nutrient-poor sites across layers; it was higher in litter but lower in soil at nutrient-poor than at nutrient-rich sites (Site condition × Depth interaction; F_2,68_ = 26.21, P < 0.001). Dried samples were grinded in a ball mill, and total carbon and nitrogen concentrations were determined by an elemental analyzer (NA 1110, CE-instruments, Rodano, Milano, Italy). Litter mass was estimated by drying at 50°C for > 48 h (Macfadyen, 1961).

### 2.3. Microbial basal respiration and biomass

Microbial biomass was measured by substrate-induced respiration with respiration measured as O_2_ consumption (Anderson and Domsch, 1978; Scheu, 1992). Samples stored at −20°C were thawed and incubated overnight at room temperature (20°C) before placing in an automated micro-respirometer. The system absorbs CO_2_ using KOH solution and the O_2_ consumed by microorganisms is replaced through electrolytic release of O_2_ from CuSO_4_ solution. Microbial basal respiration was measured as mean consumption of O_2_ during 10–23 h after attachment of the vessels to the respirometer (μg O_2_ g^−1^ h^−1^). Microbial biomass was determined after the addition of glucose to saturate microbial glycolytic enzymes based on the maximum initial respiratory response 4–7 h after the addition of D-glucose (MIRR; μl O_2_ g^−1^ h^−1^) and converted to microbial biomass (C_mic_; μg C_mic_ g^−1^) as 38 × MIRR (Beck et al., 1997). Basal respiration and substrate-induced respiration were measured at 22°C in a water bath. Fresh litter and soil from 0–5 and 5–10 cm depth equivalent to approximately 0.3, 1.0 and 3.5 g dry mass was supplemented with 80, 20 and 8 mg glucose solved in 400 μl H_2_O, respectively. The ratio between microbial basal respiration and microbial biomass was taken as microbial specific respiration (qO_2_; μg O_2_ μg^−1^ C_mic_ h^−1^).

### 2.4. Phospholipid fatty acid analysis

To quantify the composition of phospholipid fatty acids (PLFAs), lipids were extracted using a modified Bligh and Dyer method (Frostegård et al., 1993; Pollierer et al., 2015). In short, lipids were fractionated into neutral lipids, glycolipids and phospholipids by elution through silica acid columns using chloroform, acetone and methanol, respectively (0.5 g silicic acid, 3 ml; HF BOND ELUT-SI, Varian Inc., Darmstadt, Germany). Phospholipids were subjected to mild alkaline methanolysis and fatty acid methyl esters were identified by chromatographic retention time compared to standards (FAME CRM47885, C11 to C24; BAME 47080-U, C11 to C20; Sigma-Aldrich, Darmstadt, Germany) using a GC-FID Clarus 500 (PerkinElmer Corporation, Norwalk, USA) equipped with an Elite 5 column (30 m × 0.32 mm inner diameter, film thickness 0.25 μm). The temperature program started with 60°C (hold time 1 min) and increased by 30°C per min to 160°C, and then by 3°C per min to 280°C. The injection temperature was 250°C and helium was used as carrier gas. Approximately 2 g of fresh litter and 4 g of fresh soil were used for the extraction.

### 2.5. Stress indicators and fatty acid markers

The ratio of cyclopropyl PLFAs to their monoenoic precursors [cy/pre; (cy17:0 + cy19:0)/(16:1ω7 + 18:1ω7)] and the ratio of saturated to monounsaturated PLFAs [sat/mono; (14:0 + 15:0 + 16:0 + 17:0 + 18:0)/(16:1ω7 + 17:1 + 18:1ω9 + 18:1ω7)] were used as indicators of physiological or nutritional stress (Pollierer et al., 2015). The ratio of Gram^+^ to Gram^−^ bacteria was used as indicator of carbon availability (Fanin et al., 2019). The saturated fatty acids i15:0, a15:0, i16:0, i17:0 were used as markers for Gram^+^ bacteria, and the fatty acids cy17:0, cy19:0, 16:1ω7 and 18:1ω7 were assigned as markers for Gram^−^ bacteria (Zelles, 1999; Fanin et al., 2019). Bacteria were represented by the sum of Gram^+^ and Gram^−^ bacteria. Linoleic acid 18:2ω6,9 was used as fungal marker (Frostegård and Bååth, 1996). Total amount of PLFAs included all identified PLFAs (n = 40; nmol g^−1^ dry weight) and was used to calculate PLFA proportions. All stress indicators and PLFA markers were analyzed from proportions (mole percentage).

### 2.6. Statistical analyses

To estimate the effect size of Forest type and Site condition, we first fitted linear mixed models (LMMs) to log-transformed response variables and then applied planned contrasts (Piovia-Scott et al., 2019). All LMMs included Forest type (European beech, Douglas-fir, Douglas-fir/European beech, Norway spruce, Norway spruce/European beech), Site condition (nutrient-rich and nutrient-poor sites) and Depth (litter, 0–5 and 5–10 cm) as fixed effects. Models were stepwise selected by likelihood ratio test, and minimal models included all main effects and the interaction of Forest type and Site condition. The 40 forest plots were included as random effects to account for non-independence of samples from the same plot. Based on Akaike Information Criterion (AIC), the eight sites were included as random effect, and the mean-variance relationship was accounted for by a dispersion parameter to meet the assumption of homogeneity of variance (Zuur et al., 2009). Univariate response variables included microbial basal respiration, microbial biomass, microbial specific respiration, stress indicators and PLFA markers.

Contrasts for the effect size of Forest type were designed to compare coniferous and mixed forests to beech forests. European beech, the climax tree species in lowland and lower montane regions of Central Europe, was used as reference (Leuschner et al., 2017). Due to the property of log-transformed response variables, the planned contrasts are analog to log response ratios (Piovia-Scott et al., 2019). To improve interpretation, we back transformed the log response ratio into response ratio and defined it as effect size. Effect sizes of Forest type were estimated for nutrient-poor and nutrient-rich sites. In addition, we applied contrasts between nutrient-poor vs. nutrient-rich sites in a similar manner to estimate the effect sizes of Site condition.

To inspect for effects of Forest type, Site condition and their interactions on microbial community structure, we first arcsine root transformed PLFA composition, and then reduced the dimensions of fatty acids by principal component analysis at each sample depth. Multivariate analyses of variance (MANOVAs) at each sample depth were applied to principal components (PCs) to test for statistical significance. Further, selected environmental variables were included into MANOVAs as covariates to test whether they explain main effects in the models (Bennett et al., 2020). Significant PCs were determined by broken stick criterion. Covariates were selected according to permutation tests based on adjusted R^2^ in redundancy analyses (RDA). Only fatty acids ≥0.2% mean mole proportion were included in the analyses.

All analyses were done in R 4.0.3 (R Core Team, 2020). We used the ‘nlme’ package to fit LMMs and the ‘emmeans’ package to conduct planned contrasts. All mixed models met the assumptions of normality of residuals and homogeneity of variance.

## 3. Results

### 3.1. Functional indicators of microorganisms

Forest type consistently affected microbial basal respiration, microbial biomass and stress indicators at nutrient-poor but not at nutrient-rich sites (Forest type × Site condition interactions; all P < 0.08; Figs 1, 2, Table 2), and this was generally true across soil layers (Fig. S2). At nutrient-poor sites, microbial basal respiration and microbial biomass in Douglas-fir were 65% and 59% lower compared to pure beech forests. Forest type effects were similar in the two conifer-beech mixtures (−46 to −56%) and least pronounced in pure spruce forests (−33 to −36%; Fig. 1). In addition, at nutrient-poor sites microbial specific respiration was 19% lower in Douglas-fir than in beech forests.

**Figure 1.**
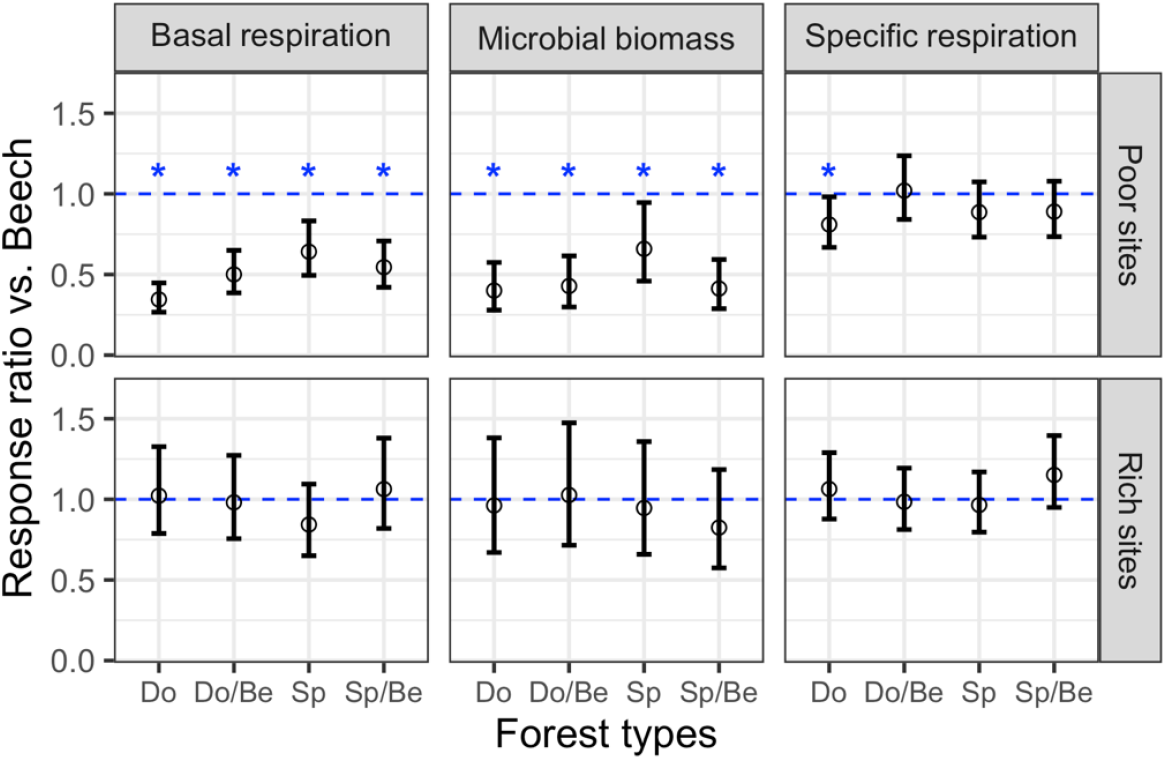
Effects of conifers and conifer-beech mixtures on microbial basal respiration (μg O_2_ g^−1^ C h^−1^), microbial biomass (μg C_mic_ g^−1^ C) and microbial specific respiration (μg O_2_ μg^−1^ C_mic_ h^−1^) at nutrient-poor and nutrient-rich sites (Douglas-fir [Do], Douglas-fir with beech [Do/Be], Norway spruce [Sp] and Norway spruce with beech [Sp/Be]). Effect sizes are given as back transformed log response ratios compared to beech forests [ln (value in coniferous or mixed forest types / values in beech)]. Effect sizes were averaged across three depths (litter, 0–5 and 5–10 cm soil). Values smaller than 1 indicate higher values in beech. Asterisks indicate significant effects (p < 0.05). Bars represent 95% confidence intervals (n=12).

**Figure 2.**
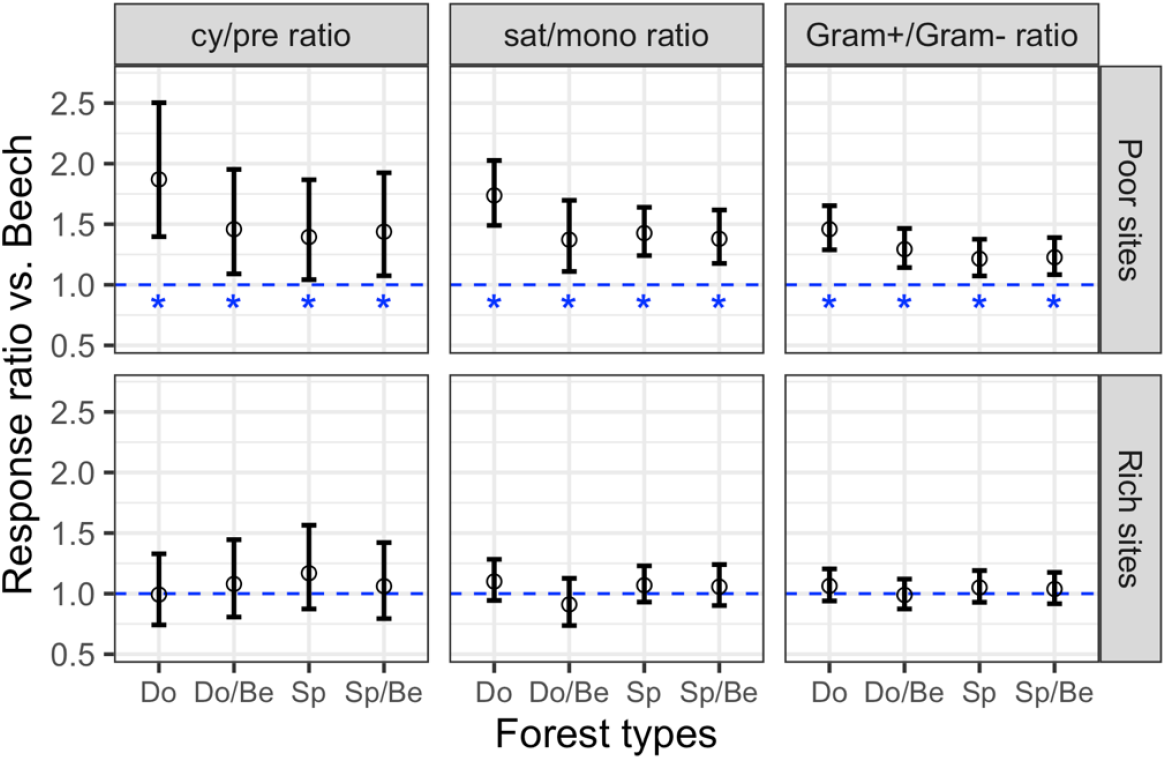
Effects of conifers and conifer-beech mixtures on stress indicators: ratio of cyclopropyl PLFAs to its monoenoic precursors (cy/pre), ratio of saturated to monounsaturated PLFAs (sat/mono), and ratio of Gram^+^ to Gram^−^ bacteria (Gram^+^/Gram^−^) at nutrient-poor and nutrient-rich sites (Douglas-fir [Do], Douglas-fir with beech [Do/Be], Norway spruce [Sp] and Norway spruce with beech [Sp/Be]). Effect sizes are given as back transformed log response ratios compared to beech forests [ln (value in coniferous or mixed forest types / values in beech)]. Effect sizes were averaged across three depths (litter, 0–5 and 5–10 cm soil). Values larger than 1 indicate lower values in beech. Asterisks indicate significant effects (p < 0.05). Bars represent 95% confidence intervals (n=12).

**Table 2.**
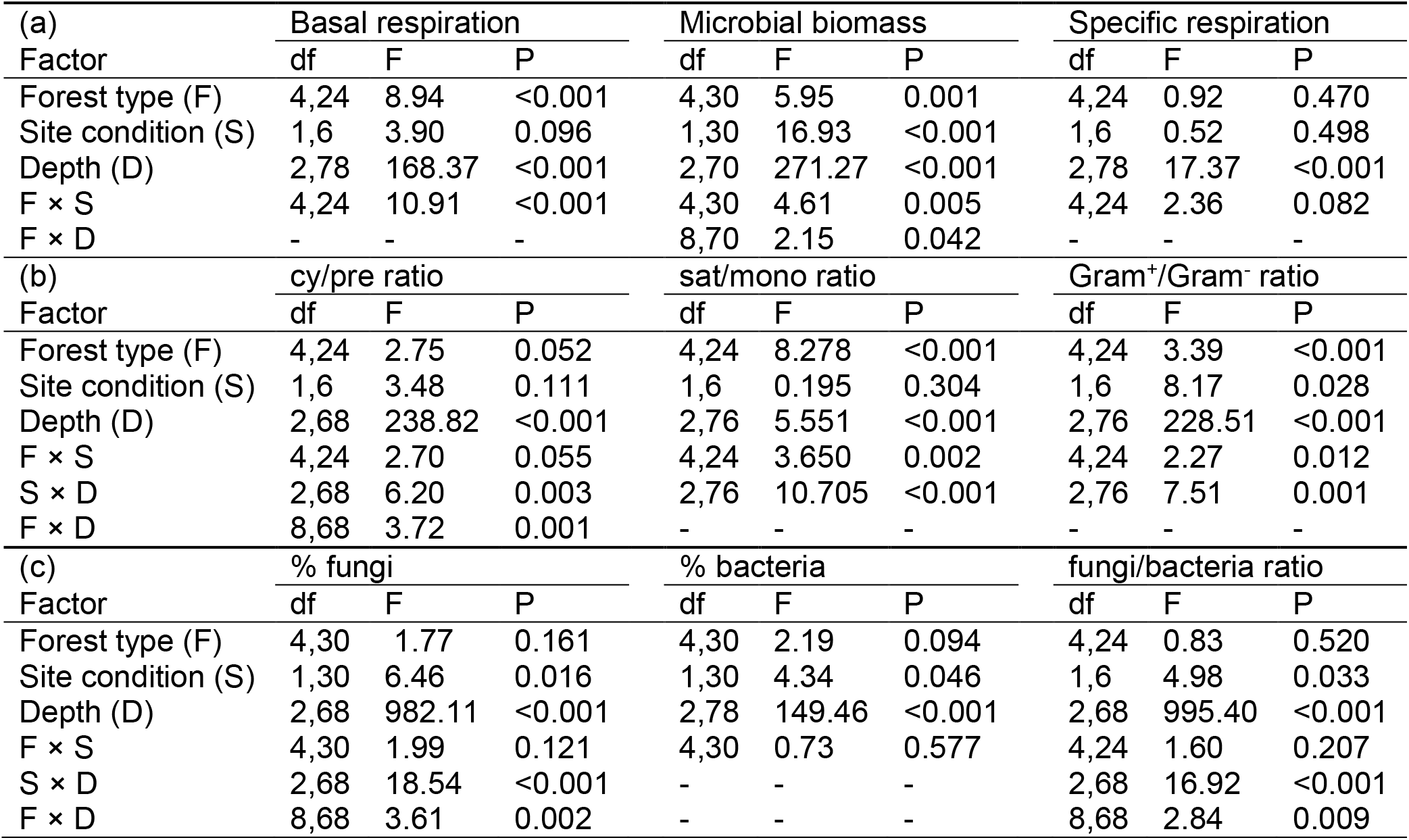
F- and P-values of linear mixed-effects models on the effect of Forest type (European beech, Douglas-fir, Norway spruce, mixture of European beech with Douglas-fir and mixture of European beech with Norway spruce), Site condition (nutrient-poor and nutrient-rich sites) and Depth (litter, 0–5 and 5–10 cm) on (a) basal respiration, microbial biomass and microbial specific respiration, (b) stress indicators (ratio of cyclopropyl PLFAs to its monoenoic precursors [cy/pre], ratio of saturated to monounsaturated PLFAs [sat/mono], Gram^+^/Gram^−^ ratio), and (c) percentages of fungal and bacterial PLFAs (of total PLFAs), and fungi/bacteria ratio. Significant effects are given in bold (p < 0.05).

At nutrient-poor sites, stress indicators, the cy/pre ratio, the sat/mono ratio as well as the Gram^+^/Gram^−^ bacteria ratio, all were lower in beech forests than in the other forest types (Fig. 2). Forest type effects on the cy/pre and sat/mono ratio were more pronounced in Douglas-fir (+87% and +74%) and less pronounced in spruce and conifer-beech mixtures (+37 to +46%). Effects of Forest type on the Gram^+^/Gram^−^ bacteria ratio also were strongest in Douglas-fir forests (+46%) and less strong in the other forest types (+21 to +29%). Stress indicators generally did not differ significantly at nutrient-rich sites (P > 0.31; Figs 2, S2, Table S1).

The response of microorganisms to changes in site conditions also varied with forest types. Differences between nutrient-rich and nutrient-poor sites were largest in Douglas-fir and smallest in beech forests (Figs 3, 4, Table S2). At nutrient-poor sites, microbial basal respiration in Douglas-fir forests was 57% lower and microbial biomass in pure and mixed forests of Douglas-fir both were 47% lower compared to nutrient-rich sites. In parallel, but less strong, microbial biomass in spruce mixed forests was 36% lower at nutrient-poor than that at nutrient-rich sites.

**Figure 3.**
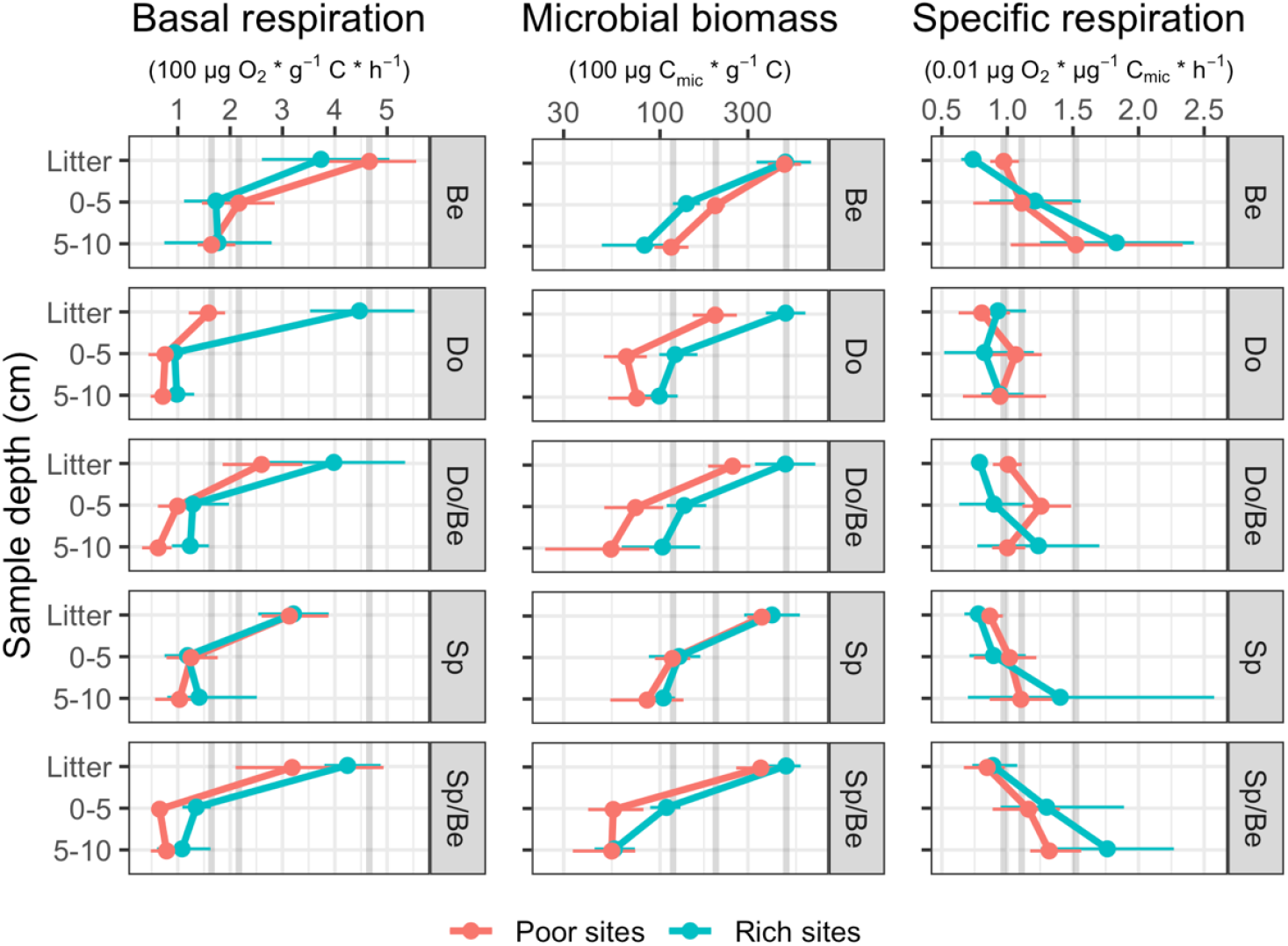
Changes in microbial basal respiration (μg O_2_ g^−1^ C h^−1^), microbial biomass (μg C_mic_ g^−1^ C) and microbial specific respiration (μg O_2_ μg^−1^ C_mic_ h^−1^) with soil depth (litter and 0–5 and 5–10 cm soil) in five forest types (European beech [Be], Douglas-fir [Do], Douglas-fir with beech [Do/Be], Norway spruce [Sp] and Norway spruce with beech [Sp/Be]) at nutrient-poor and nutrient-rich sites. Points and horizontal bars represent means and standard errors (n=4). The grey vertical bars represent respective values in beech forests at nutrient-poor sites in litter, 0–5 and 5–10 cm soil. Note log scale for microbial biomass.

**Figure 4.**
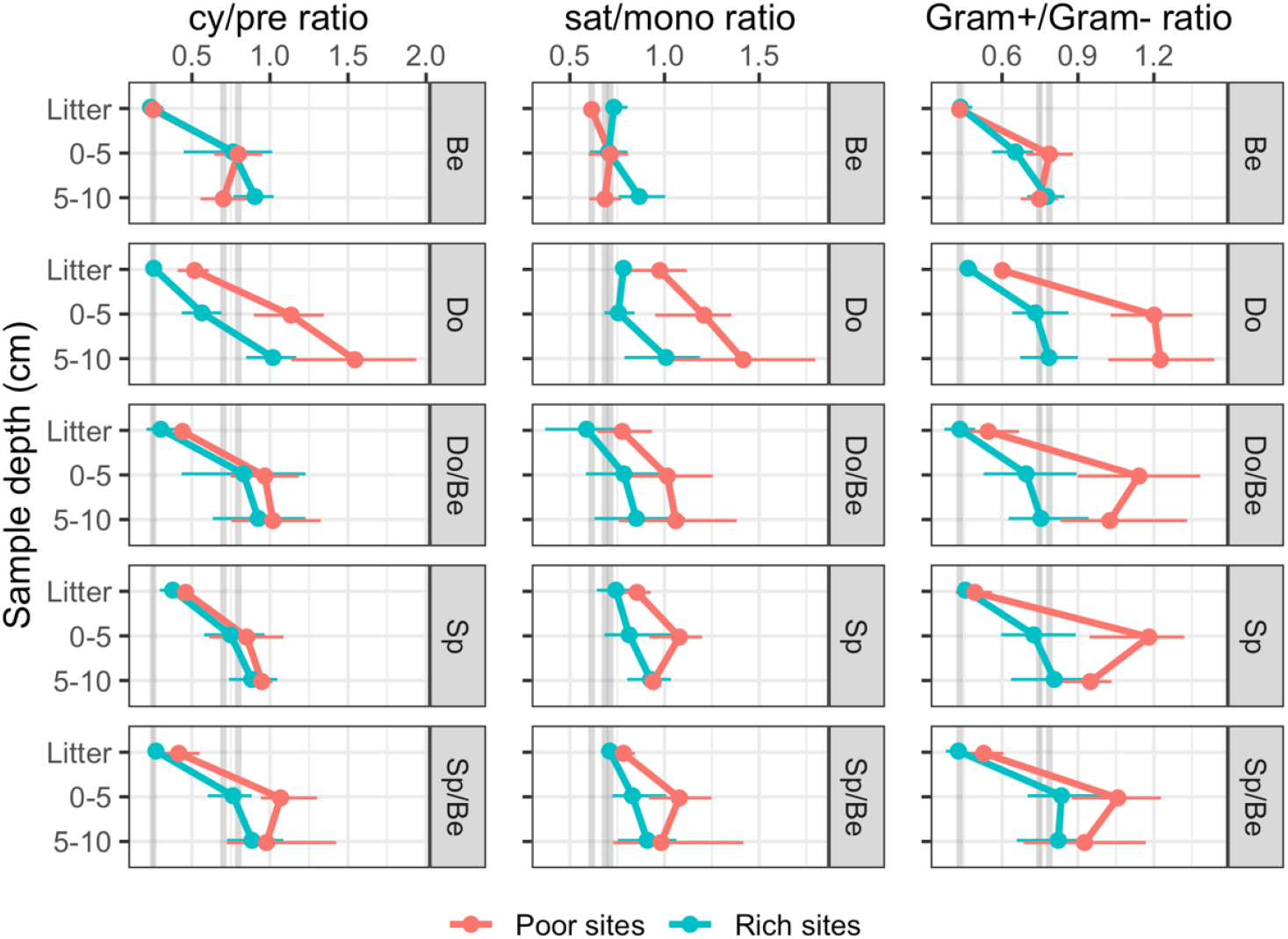
Changes in stress indicators (ratio of cyclopropyl fatty acids to its precursors [cy/pre], ratio of saturated to monounsaturated fatty acids [sat/mono] and ratio of Gram^+^ to Gram^−^ bacteria [Gram^+^/Gram^−^]) with soil depth (litter and 0–5 and 5–10 cm soil) in five forest types (European beech [Be], Douglas-fir [Do], Douglas-fir with beech [Do/Be], Norway spruce [Sp] and Norway spruce with beech [Sp/Be]) at nutrient-poor and nutrient-rich sites. Points and horizontal bars represent means and standard errors (n=4). The grey vertical bars represent respective values in beech forests at nutrient-poor sites in litter, 0–5 and 5–10 cm soil.

Likewise, in Douglas-fir forests the stress indicator cy/pre ratio at nutrient-poor sites exceeded that at nutrient-rich sites by 82%, and similarly, the sat/mono and Gram^+^/Gram^−^ bacteria ratio at nutrient-poor sites exceeded that at nutrient-rich sites by 38–40% (Fig. 4, Table S2). In Douglas-fir mixed forests the Gram^+^/Gram^−^ bacteria ratio at nutrient-poor sites exceeded that at nutrient-rich sites by 34%. Further, the response of microbial stress to changes in site conditions varied with soil depth. In particular, the responses of the sat/mono ratio and the Gram^+^/Gram^−^ ratio to site conditions were most pronounced at 0–5 cm soil depth.

### 3.2. Microbial community structure

As indicated by the PLFA patterns, microbial community composition in beech forests differed from that in Norway spruce and Douglas-fir forests, with mixed forests being intermediate between pure beech and coniferous forests (Fig. 5). Across layers, Forest type effects on microbial community composition were significant in litter and 0–5 cm soil, and marginally significant in 5–10 cm soil (Table 3). Despite strong turnover of microbial community composition between nutrient-rich and nutrient-poor sites (Site condition effects; all p < 0.001), in beech forests microbial community structure in nutrient-poor and nutrient-rich sites was similar, and this was most apparent in 0–5 cm soil (Forest type × Site condition interaction; Fig. 5, Table 3).

**Table 3.**
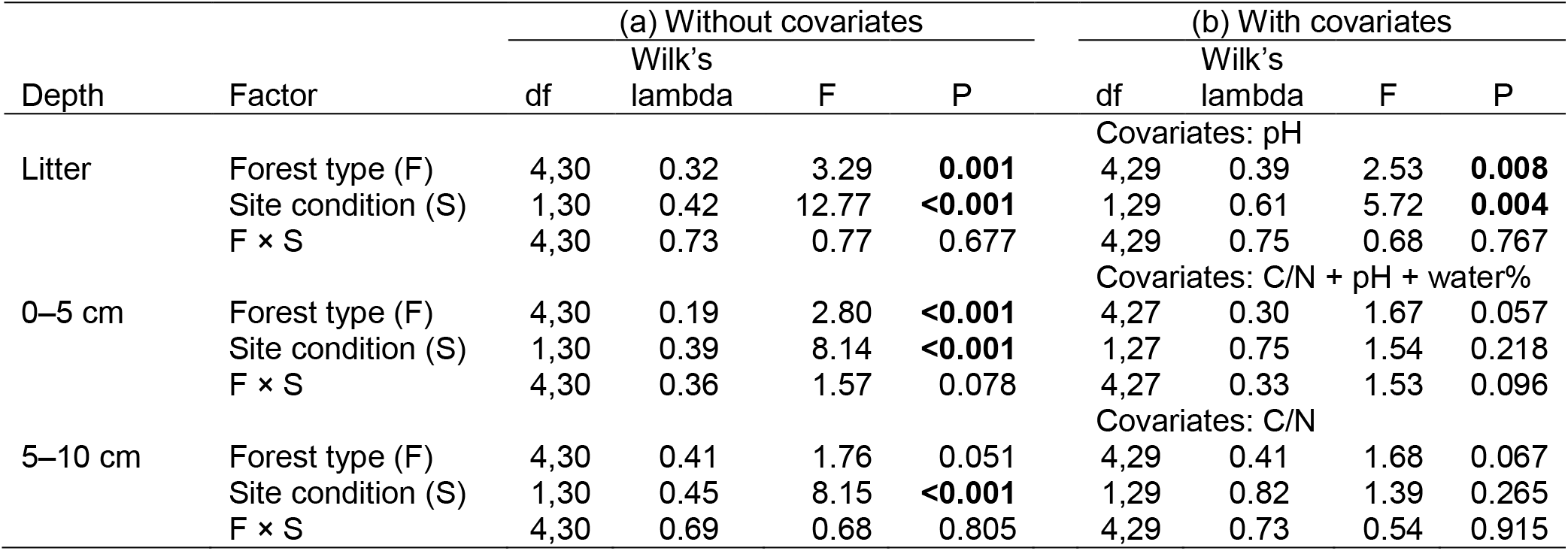
Results of two-way MANOVAs on PLFA composition (a) without covariates and (b) with covariates (pH, C/N ratio, water content) in litter, 0–5 and 5–10 cm soil. Factors include Forest type (European beech, Douglas-fir, Norway spruce and mixture of European beech with Douglas-fir and mixture of European beech with Norway spruce), Site condition (nutrient-rich and nutrient-poor sites) and their interactions. MANOVAs were based on principal components from each sample depth (significant PCs were determined by broken stick criterion; see Methods). Type I sequential sum of square was used for models with and without covariates. Significant effects are given in bold (P < 0.05).

**Figure 5.**
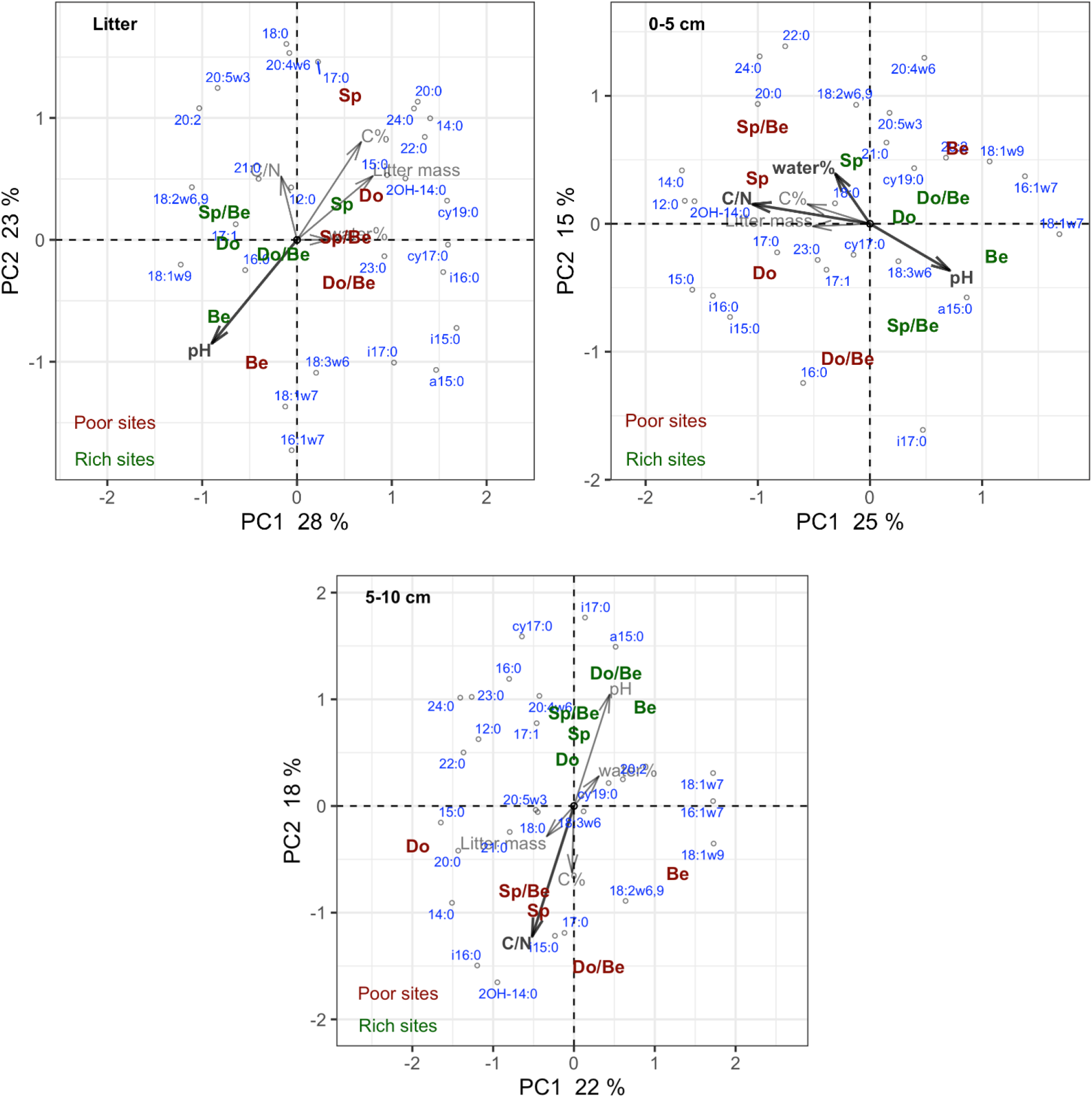
Principal component analysis of the microbial community structure (as indicated by phospholipid-derived fatty acids) in five forest types (European beech [Be], Douglas-fir [Do], Douglas-fir with beech [Do/Be], Norway spruce [Sp] and Norway spruce with beech [Sp/Be]) at nutrient-poor and nutrient-rich sites in litter, 0–5 and 5–10 cm soil depth. Positions of forest types represent centroids (n=4). Nutrient-poor sites are given in red and nutrient-rich sites in green. C/N, pH, water content (water%), carbon content (C%) as well as litter mass were post-fitted into the plot. Significant variables selected by RDA permutation test were in bold (based adjusted R^2^).

A number of fatty acids varied among forest types and with site conditions. Beech forests were characterized by unsaturated fatty acids at all depths, in particular by 16:1ω7, 18:1ω7, 18:1ω9 (Fig. 5). Norway spruce forests were associated with long-chain saturated fatty acids (22:0, 24:0), and Douglas-fir forests with bacterial marker fatty acids (2OH-14:0, cy17:0). Further, nutrient-rich sites were associated with unsaturated fatty acids in litter, such as 18:1ω9 and 18:2ω6,9.

Litter at nutrient-rich sites was generally richer in fungi than at nutrient-poor sites, but the opposite was true for the soil (Site condition × Depth interaction; Fig. S3, Table 2). The relative abundance of fungi and bacteria generally did not differ among forest types, but the relative abundance of fungi consistently decreased, whereas the relative abundance of bacteria increased with soil depth (Figs S3, S4). In addition, based on the contrast of Site condition, at nutrient-poor sites the relative abundance of fungi in pure beech and spruce forests was higher than at nutrient-rich sites. By contrast, in pure and mixed Douglas-fir forests at nutrient-poor sites the relative abundance of fungi in soil was similar to that at nutrient-rich sites (Fig. S3, Table S2).

Litter mass did not differ significantly among forest types, but was 58% higher at nutrient-poor than at nutrient-rich sites (Forest type: F_4,30_ = 0.55, P = 0.70; Site condition: F_1,30_ = 13.21, P = 0.001; Fig. S5). Nutrient-poor sites also were more nitrogen-limited and acidic than nutrient-rich sites (Fig. 5). Soil C/N ratio, pH and soil moisture were among the most important environmental variables in explaining variations in PLFA composition (Table S3). In litter, pH accounted for most (8%) of the variation explained by all environmental variables (13%). By contrast, in 0–5 cm as well as 5–10 cm soil depth, C/N ratio accounted for most of the variation explained by all environmental variables (22% out of the total of 37% and 13% out of the total of 15%, respectively; Table S3). Underlining the importance of environmental factors in structuring microbial communities at the study sites, the main effect of Site condition became non-significant in soil after including C/N ratio, pH, water content (0–5 cm soil depth) and C/N ratio (5–10 cm soil depth) as covariates in MANOVAs. By contrast, in litter, including pH as covariate did not significantly affect the effects of Site condition. The effects of Forest type remained marginally significant in both depths of soil and significant in litter when including covariates (Fig. 5, Table 3).

## 4. Discussion

### 4.1. Functional indicators

The present study evaluated effects of two coniferous trees in pure and mixed stands on soil microbial community composition and functioning covering a range of nutrient conditions, with native European beech as reference. Overall, the results in part support our first and second hypotheses. Forest type only affected the functioning of microbial communities at nutrient-poor sites, but not at nutrient-rich sites. At nutrient-poor sites, basal respiration, microbial biomass and stress indicators all were detrimentally affected in pure and mixed coniferous forests, and, contrasting our first hypothesis the effects differed between Douglas-fir and spruce forests, with the effects being particularly strong in Douglas-fir, but less pronounced in Norway spruce and mixed stands. Enrichment of beech with either Douglas-fir or spruce in mixed forests both compromised microbial functioning, suggesting that caution is needed when admixing conifers to European beech forests at nutrient-poor sites. By contrast, similar basal respiration, microbial biomass and stress indicators at nutrient-rich sites across the studied forest types contradict our second hypothesis and suggest that at nutrient-rich sites soil microbial communities are rather irresponsive to changes in tree species. This implies that at nutrient-rich sites planting Douglas-fir in pure or mixed forests may provide an alternative to planting Norway spruce.

The site- and tree species-specific responses of microorganisms are likely caused by differences in the provisioning of carbon resources. Gram^+^ bacteria better cope with recalcitrant carbon resources, whereas Gram^−^ bacteria favor labile carbon compounds (Kramer and Gleixner, 2008; Fanin et al., 2019). High Gram^+^/Gram^−^ ratio in Douglas-fir and low Gram^+^/Gram^−^ ratio in European beech at nutrient-poor sites therefore indicate more pronounced shortage of labile carbon resources in Douglas-fir compared to beech forests. This is further supported by the lower microbial specific respiration in Douglas-fir compared to beech forests, which was largely due to lower microbial basal respiration (Anderson and Domsch, 2010). Low availability of labile carbon favors oligotrophic microbial communities characterized by low respiration rates and biomass (Potthast et al., 2010; Fanin et al., 2019), and this is consistent with our findings of low basal respiration and microbial biomass in pure and mixed coniferous forests at nutrient-poor sites.

Moreover, higher ratio of cyclopropyl PLFAs to their monoenoic precursors and higher ratio of saturated to monounsaturated PLFAs in pure and mixed coniferous compared to pure beech forests at nutrient-poor sites indicates that microorganisms in pure and mixed coniferous forests were more nutrient (and/or water) stressed (Moore-Kucera and Dick, 2008; Pollierer et al., 2015). More stressed microorganisms in coniferous and mixed compared to pure beech forests at nutrient-poor sites also is indicated by lower proportions of the presumably Gram^−^ bacteria marker fatty acids 16:1ω7 and 18:1ω7 in the former. Under stress, Gram^−^ bacteria change the composition of their cell membranes from 16:1ω7 and 18:1ω7 to cyclopropane fatty acids associated with slower growth rates (Kieft et al., 1994; Lundquist et al., 1999). The similarity of microbial indicators in beech forests at nutrient-rich and nutrient-poor sites indicates that, compared to coniferous stands, beech maintained microbial community functioning and prevented stress conditions for microorganisms by providing more labile carbon resources at nutrient-poor conditions, presumably by releasing high amounts of root-derived resources (Meier et al., 2020).

The suggestion that beech is able to alleviate microbial stress via maintaining or even increasing the release of root-derived resources at nutrient-poor sites is supported by higher fine root biomass in beech than in pure and mixed coniferous forests at nutrient-poor sites, whereas at nutrient-rich sites fine root biomass varies little among forest types (A. Lwila, pers. comm.). If soil nutrient availability declines, plants allocate surplus carbon into roots, associated with increased root exudation (Laliberté et al., 2017; Prescott et al., 2020). Recently, higher root exudation has been confirmed for beech forests at more acidic and nitrogen deficient stands (Meier et al., 2020), conditions similar to our nutrient-poor sites characterized by higher C/N ratio and more acidic soil than the nutrient-rich sites. High amounts of root-derived resources presumably also were responsible for the alleviated microbial resource deficiency and stress in beech forests at nutrient-poor sites. Trenching and girdling experiments also confirmed that reducing or omitting the flux of root-derived resources into the soil strongly reduces fungal and bacterial biomass, demonstrating the importance of root-derived resources for maintaining soil microbial biomass (Kaiser et al., 2010; Bluhm et al., 2019). Thus, beech may facilitate soil microorganisms at environmental stress conditions by increasing root exudation contrasting both of the studied coniferous species. This may also mitigate detrimental effects of Douglas-fir on microorganisms at nutrient-poor sites in mixed forests. Generally, high amounts of root-derived resources in beech may be responsible for the facilitation of neighboring coniferous trees by beech, resulting in overyielding in mixed stands at nutrient-poor sites (Toïgo et al., 2015; Ammer, 2019; David et al., 2020).

Notably, we sampled litter and soil during autumn / winter after beech had shed its leaves, which might have affected microbial communities. However, any litter effect on microbial communities should have been similar in nutrient-rich and nutrient-poor sites, as litter input in beech forests varies little with site conditions (Meier et al., 2005). By contrast, we found microbial community characteristics to only differ among forest types at nutrient-poor sites, and litter mass on the forest floor did not differ among forest types (Fig. S5). Supporting our suggestion on the importance of root-derived resources, at nutrient-poor sites fine root biomass in beech forests exceeded that in coniferous forests by more than 1.6 times, whereas fine root biomass did not differ significantly among forest types at nutrient-rich sites (A. Lwila, pers. comm.). The hypothesis of high root-derived resources in beech forests implies that the impact of roots on microorganisms still was strong in autumn / winter indicating that not only recent photosynthates but also stored carbohydrates contribute to providing root-derived resources to soil microorganisms (Druebert et al., 2009). However, it remains open whether the effects of Forest type on microbial communities get stronger or weaker with time, hampering generalization of our finding across seasons. Although microbial communities fluctuate considerably with season (Koranda et al., 2013; Abramoff and Finzi, 2016; Nacke et al., 2016), comprehensive sampling in Douglas-fir forests across seasons suggests that, except during summer drought, microbial stress indicators fluctuate little (Moore-Kucera and Dick, 2008). Further, although fine roots generally are concentrated in upper soil layers, we may not have captured the full effects of roots, as our sampling was limited to the upper 10 cm of the soil, but e.g., beech roots may extend considerably deeper into the soil (Leuschner et al., 2004). Experimental manipulations of root-derived resources in the field are needed to fully resolve the role of root-derived resources in driving microbial community structure and functioning (Koranda et al., 2011; Bluhm et al., 2019).

Contrary to our third hypothesis, the effects of Forest type were not stronger in litter than in soil. Forest types impacted microbial respiration, biomass and stress indicators similarly across layers at nutrient-poor sites. This uniformity further supports our conclusion that effects of Forest type were not the result of different litter quantity and quality, but were due to root-derived resources. This contrasts earlier suggestions that, because litter is not buffered against environmental conditions, microbial responses in litter are more sensitive to environmental change than those in soil (Pollierer et al., 2015). Reflecting environmental changes, microbial biomass decreased while microbial stress indicators increased with soil depth. This sensitivity may be related to lower carbon availability deeper in soil, resulting in stronger resource limitation (Fierer et al., 2003). As indicated by Gram^+^/Gram^−^ ratios, the difference in resource limitation between litter and soil was greatest in pure coniferous and mixed forests at nutrient-poor sites.

### 4.2. Community structure

We assumed that microbial functioning ultimately relies on its community structure, and this is supported by the consistent turnover in microbial community composition as well as the differences in microbial functioning between nutrient-rich and nutrient-poor sites. Despite increased resource limitation with soil depth, microbial community structure in 5–10 cm soil differed little between forest types at nutrient-rich sites. This suggests that the influence of forest types fades deeper in soil. By contrast, at nutrient-poor sites, microbial community structure differed between beech and coniferous forests in soil, particularly in 5–10 cm depth, supporting our conclusion that effects of tree species on microorganisms in nutrient-poor soil are predominantly due to differences in root-derived resources. Unlike Douglas-fir and spruce, European beech maintained and stabilized microbial community structure and functioning in litter and soil irrespective of site conditions. Interestingly, in 5–10 cm depth at nutrient-poor sites, microbial community structure and carbon limitation under Douglas-fir differed most strongly from beech. This implies that, among the tree species studied, root-derived resources are most limited under Douglas-fir at nutrient-poor sites, resulting in pronounced resource shortage and marked changes in microbial community composition. Such changes in microbial community structure may significantly impact the functioning of soil microbial communities as the availability of carbon resources and carbon limitation affect the vertical distribution of saprotrophic fungi and decomposition processes e.g., by shifting the competition between saprotrophic and mycorrhizal fungi (Gadgil and Gadgil, 1971; Lindahl et al., 2007).

Our results support previous findings that regional factors more strongly shape microbial community structure than tree species (Richter et al., 2018). The relative abundance of fungi and bacteria as well as the fungi/bacteria ratio did not vary with forest types but rather with site conditions. The relative abundance of fungi was favored at nutrient-poor sites, supporting earlier suggestions that the fungal energy channel dominates in low nutrient systems (Wardle et al., 2004). However, at nutrient-poor sites, the fungal energy channel did not dominate in Douglas-fir forests, despite high C/N ratio and low pH. In fact, the relative abundance of fungi in Douglas-fir forest soil was similar at nutrient-rich and nutrient-poor sites, resulting in lower fungal abundance in Douglas-fir than beech forests, particularly in 5–10 cm depth. In addition to low amounts of root-derived resources, low fungal abundance in Douglas-fir forest soil may be due to a mismatch between local and home (native range of Douglas-fir) fungal communities, as only a subset of native-range fungal species has been recorded in the soils to which Douglas-fir has been introduced (Schmid et al., 2014). Although molecular techniques are needed to identify the microbial taxa responsible for differences among forest types, particularly between Douglas-fir and beech at nutrient-poor sites, our study supports the sensitivity of PLFA analysis in detecting changes in microbial community composition (Ramsey et al., 2006).

Forest type effects on microbial community composition were marginally significant after including covariates, indicating that forest types affected microbial community composition independently of the studied environmental variables. By contrast, the measured environmental variables well captured differences in soils between nutrient-poor and nutrient-rich sites, and explained much of the variance in site conditions. Among the environmental variables studied, C/N ratio and pH best explained variations in microbial community structure. Soil pH typically correlates with soil nutrient availability (Meier et al., 2020), further supporting the importance of nutrients in driving microbial responses to variations in site conditions. However, factors other than soil nutrient status and soil pH may have contributed to the observed differences between nutrient-rich and nutrient-poor sites. Despite inconsistent effects of site conditions on water content across layers, and little effects of water content on litter and soil microbial community structure, nutrient-poor sites also are characterized by lower precipitation. This is supported by higher δ^13^C values of litter at nutrient-poor than at nutrient-rich sites (J.-Z. Lu, unpubl. data), reflecting higher water stress at nutrient-poor than at nutrient-rich sites (Peuke et al., 2006). Presumably, in addition to nutrients and pH, water stress contributed to the observed differences in microbial community composition and functioning between nutrient-poor and nutrient-rich sites. This is supported by the fact that soil water conditions are intricately linked to soil nutrient dynamics and the uptake of nutrients by plants (de Vries et al., 2019). To disentangle pathways linking forest stands and soil microbial communities, it is most promising to focus on water deficient, low pH and nutrient-poor stands.

### 4.3. Management implications

Due to increasing drought and associated outbreaks of bark beetles, Norway spruce forests are increasingly threatened in lowland regions of Europe. Facing such challenge, mixed forests and planting alternative tree species such Douglas-fir provide promising options for provisioning long-term ecosystem services. By studying microbial communities in pure and mixed forests covering a range of nutrient and water conditions, we found that effects of forest types on soil microorganisms vary among tree species and mixed stands, but this strongly depends on the nutrient status of the sites. At nutrient-poor sites, Douglas-fir strongly impacted microbial community structure and functioning compared to beech. Therefore, in the long-term planting non-native tree species such as Douglas-fir may compromise ecosystem functioning at nutrient-poor sites. This may not only question the establishment of pure Douglas-fir forests, but also the enrichment of beech forests by Douglas-fir at nutrient-poor sites. However, as planting Douglas-fir in single species and mixed stands with beech hardly impacted the structure and functioning of soil microbial communities at nutrient-rich sites, planting Douglas-fir may provide an alternative to Norway spruce at least at nutrient-rich sites. Although more information on ecosystem properties and processes is needed to gain a holistic understanding on the consequence of forest plantation on ecosystem functioning, we conclude that microbial stress is intensified in nutrient-poor soils by planting Douglas-fir.

## Acknowledgements

This work was supported by the German Research Foundation through the Research Training Group 2300: “Enrichment of European beech forests with conifers: impacts of functional traits on ecosystem functioning” [316045089]. We are indebted to C. Bluhm, T. Volovei, G. Humpert, M. Unger, J. Meyer and S.-X. Zhou for assistance. We thank C. Ammer, H. Riebl, T. Hartke, A. Polle, X.-M. Lu, N. Lamersdorf, Z.-P. Li, S. Bluhm, G. Schulz, T.-W. Chen, M. Pollierer, A. Lwila, K. Schumann, I. Bebre and A. Rivera for helpful discussions. Special thanks to S. Müller for coordinating the project.

