## supplemental figures and tables for "Response of soil microbial communities to mixed forests of European beech and conifers: Variations with site conditions"

### Supporting Information

<u>Supplementary Figures</u>

**Figure S1** Study sites

**Figure S2** Depth-specific effects of forest type on respiration and stress indicators across site conditions

**Figure S3** Depth-specific difference of microbial markers between nutrient-rich and nutrient-poor sites

**Figure S4** Depth-specific effects of forest type on microbial markers at nutrient-rich and nutrient-poor sites

**Figure S5** Comparison of litter mass among forest types at nutrient-rich and nutrient-poor sites

<u>Supplementary Tables</u>

**Table S1** Statistics contrasts of the effects of forest type

**Table S2** Statistics contrasts of the effects of site condition

**Table S3** Permutation tests of covariates in RDA

Figure S1

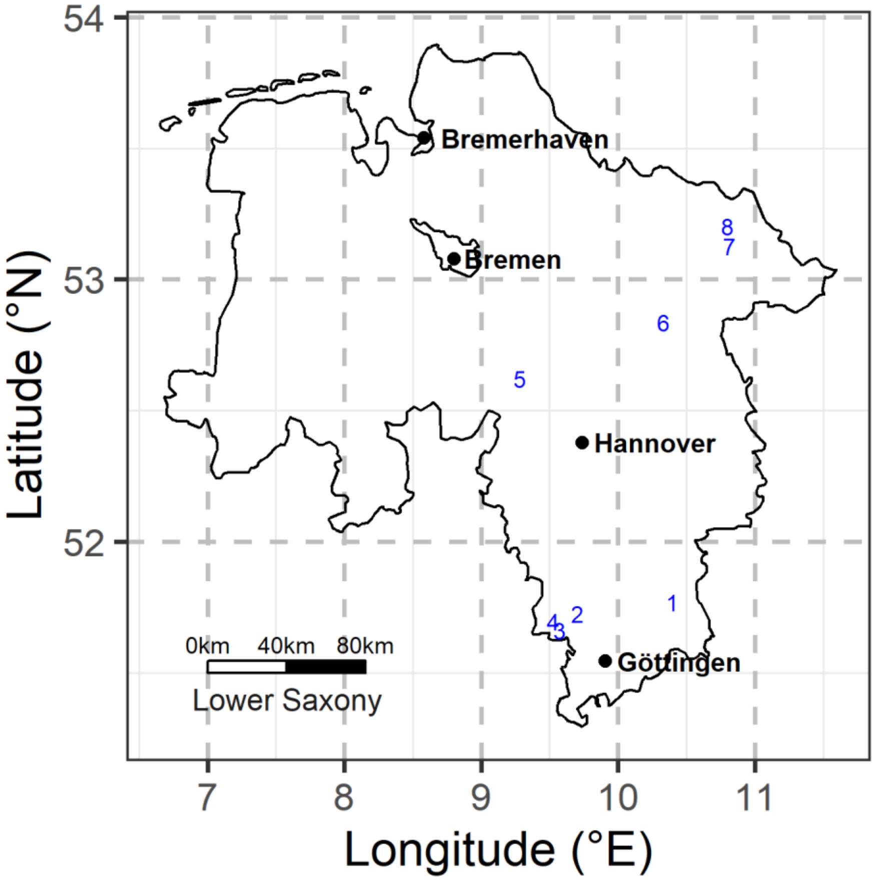


Figure S1. Study sites (number 1 to 8; see Table 1 for details). Four nutrient-rich sites and four-nutrient poor sites located in the south and north of Lower Saxony, Germany.

Figure S2

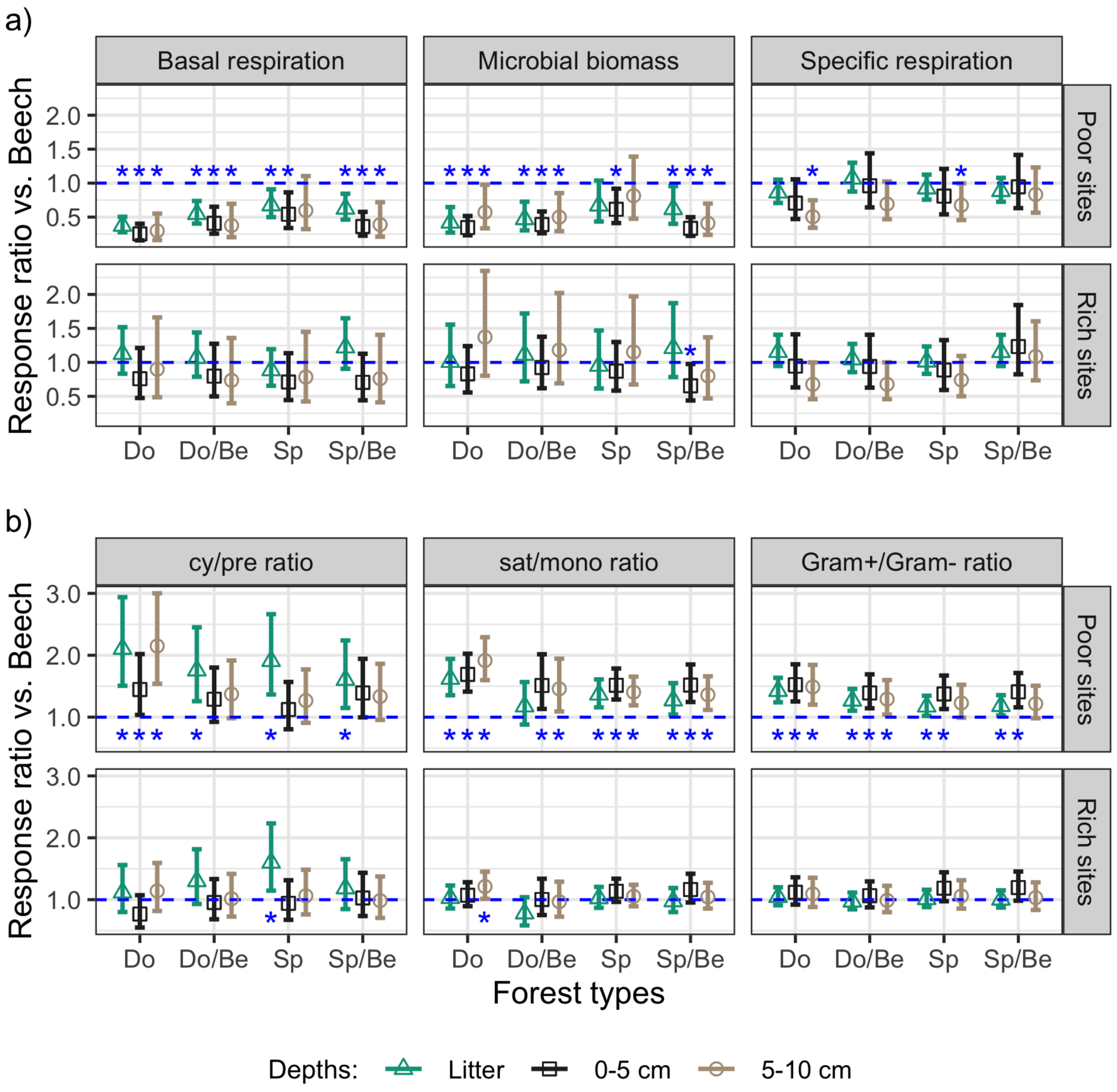


Fig. S2. (a) Depth-specific effects of conifers and conifer-beech mixtures (Douglas-fir [Do], Douglas-fir with European beech [Do/Be], Norway spruce [Sp] and Norway spruce with European beech [Sp/Be]) on microbial basal respiration (μg $O_2$ $g^{-1}$ C $h^{-1}$), microbial biomass (μg $C_{mic}$ $g^{-1}$ C) and microbial specific respiration (μg $O_2$ $μg^{-1}$ $C_{mic}$ $h^{-1}$) at nutrient-poor and nutrient-rich sites; (b) depth-specific effects on the ratio of cyclopropyl PLFAs to its monoenoic precursors, saturated to monounsaturated PLFAs, and Gram positive to Gram negative bacteria ratio (cy/pre, sat/mono and Gram+/Gram-, respectively) at nutrient-poor and nutrient-rich sites. Effect sizes are given as back transformed log response ratios compared to beech forests [ln (value in coniferous or mixed forest types / values in beech)]. Effect sizes were estimated from litter, 0–5 and 5–10 cm soil depth based on mixed-effects models with all two-way interactions. Asterisks indicate significant effects ($p < 0.05$). Bars represent 95% confidence intervals (n=4).

Figure S3

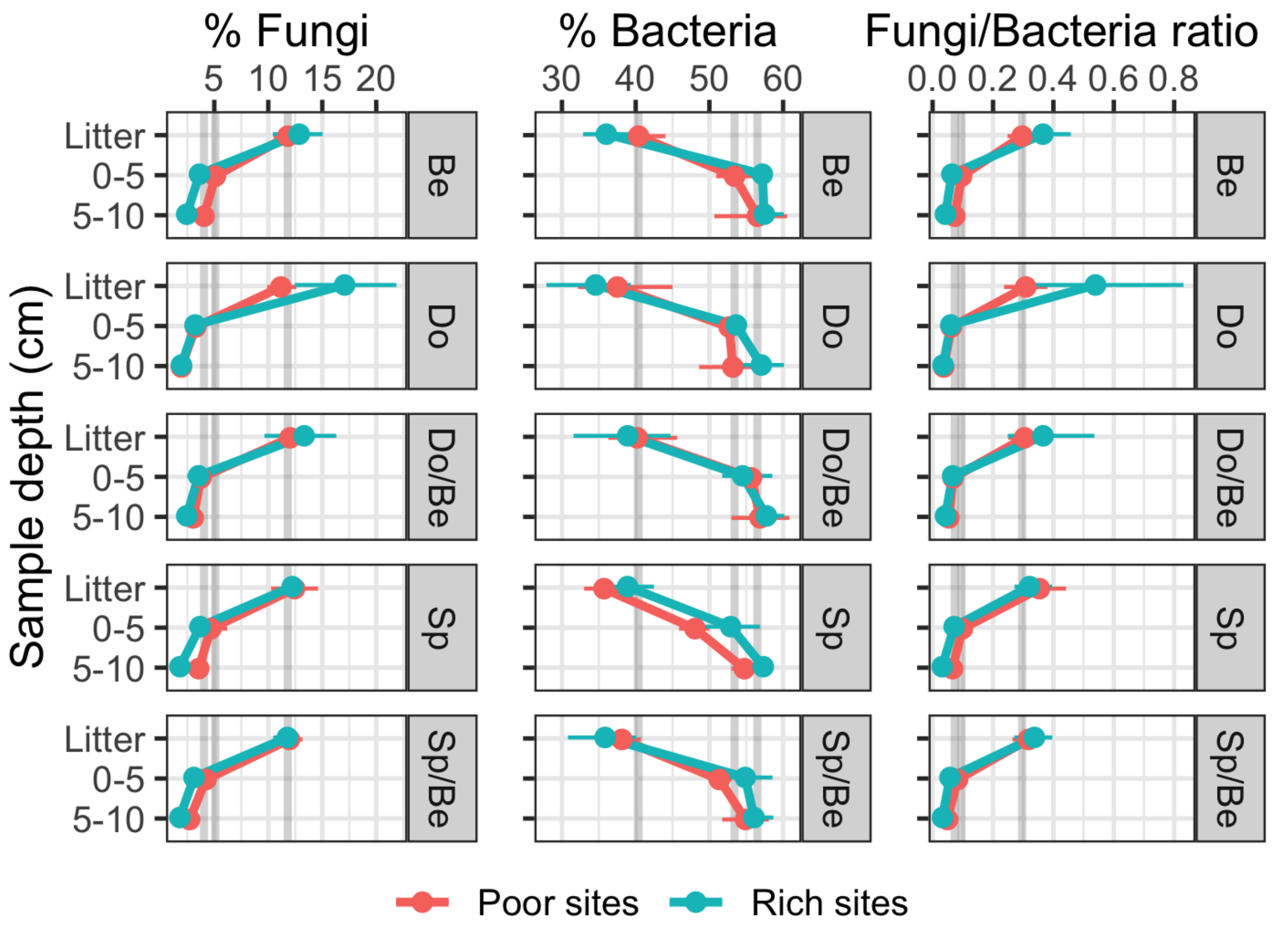


Figure S3. Changes in the relative abundance of fungi, bacteria and fungi/bacteria ratio with soil depth (litter and 0–5 cm and 5–10 cm soil) in five forest types (European beech [Be], Douglas-fir [Do], Douglas-fir with beech [Do/Be], Norway spruce [Sp] and Norway spruce with beech [Sp/Be]) at nutrient-poor and nutrient-rich sites. Points and horizontal bars represent means and standard errors (n=4). The grey vertical bars indicate respective values in beech forests at nutrient-poor sites in litter, 0–5 and 5–10 cm depth.

Figure S4

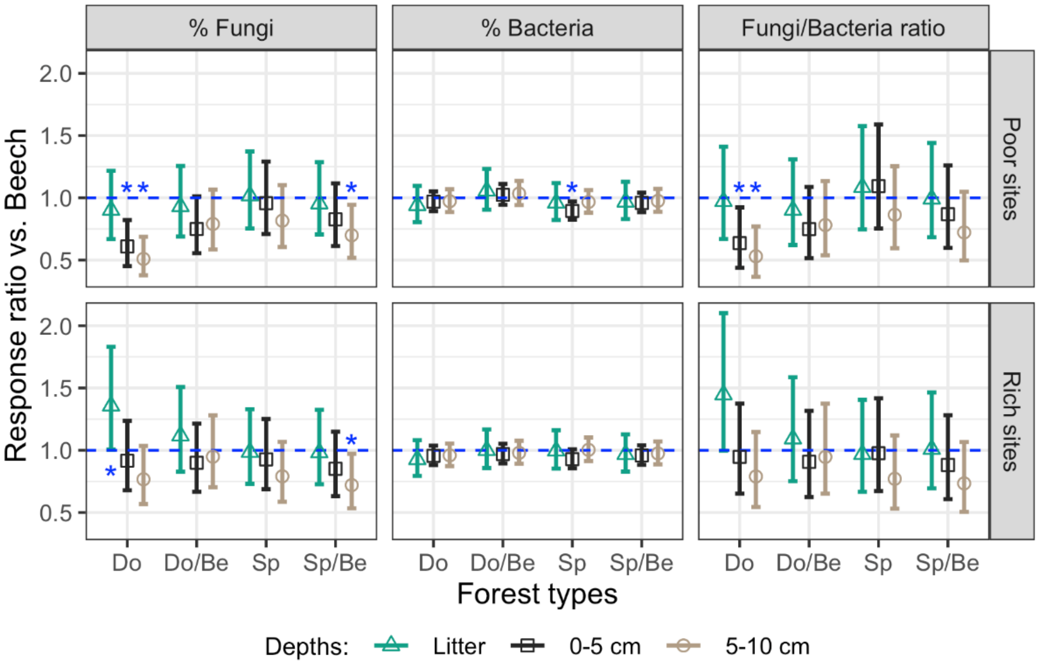


Figure S4. Depth-specific effects of conifers and conifer-beech mixtures (Douglas-fir [Do], Douglas-fir with beech [Do/Be], Norway spruce [Sp] and Norway spruce with beech [Sp/Be]) on the relative abundance of fungi, bacteria and fungi/bacteria ratio at nutrient-poor and nutrient-rich sites. Effect sizes are given as back transformed log response ratios compared to beech forests [ln (value in coniferous or mixed forest types / values in beech)]. Effect sizes were estimated from litter, 0–5 and 5–10 cm soil depth based on mixed-effects models with all two-way interactions. Asterisks indicate significant effects ($p < 0.05$). Bars represent 95% confidence intervals (n=4).

Figure S5

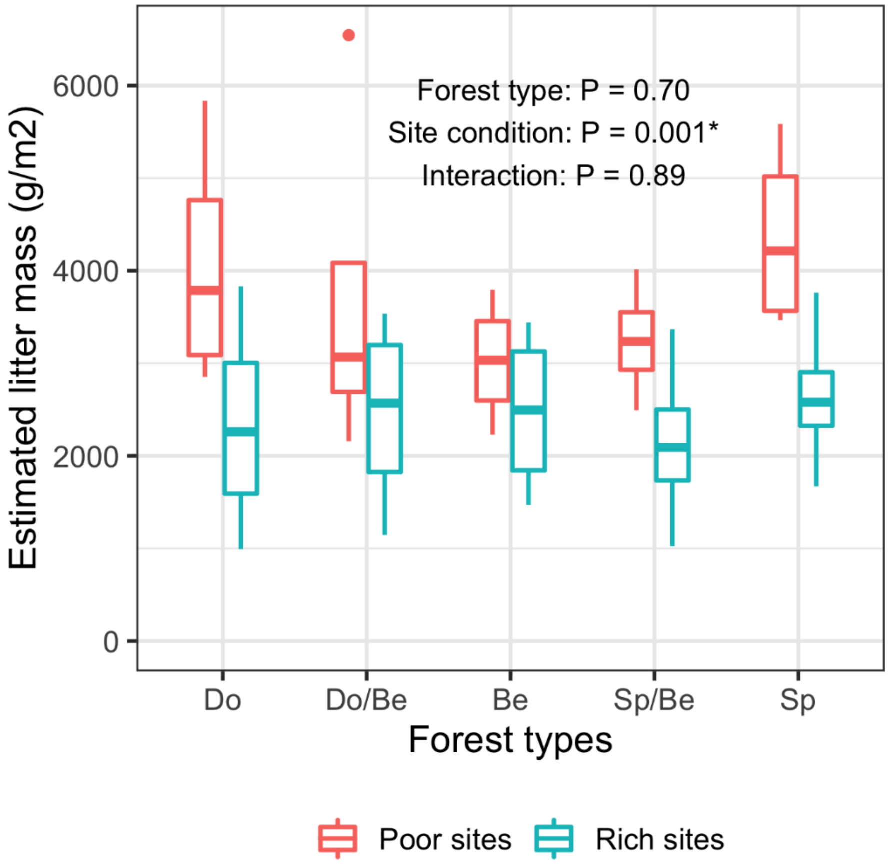


Figure S5. Litter mass in the studied forest types (Douglas-fir [Do], Douglas-fir with beech [Do/Be], European beech [Be], Norway spruce with beech [Sp/Be] and Norway spruce [Sp]) at nutrient-poor and nutrient-rich sites. Litter mass includes leaves, needles and reproductive organs on the forest floor.

Table S1. Contrasts comparing the studied forest types (Douglas-fir [Do], Douglas-fir and European beech mixture [Do/Be], Norway spruce and European beech mixture [Sp/Be]) to European beech forests (Be). Ratio was back-transformed from log response ratio. Significant p-values are given in bold ($p < 0.05$).

| **Variable** | **Contrast** | **Site** | **Ratio** | **p-value** |
|---|---|---|---|---|
| Basal respiration | Do-Be | Nutrient-rich | 1.022 | 0.862 |
| Basal respiration | Do/Be-Be | Nutrient-rich | 0.981 | 0.880 |
| Basal respiration | Sp/Be-Be | Nutrient-rich | 1.063 | 0.632 |
| Basal respiration | Sp-Be | Nutrient-rich | 0.843 | 0.190 |
| Basal respiration | Do-Be | Nutrient-poor | 0.345 | **<0.001** |
| Basal respiration | Do/Be-Be | Nutrient-poor | 0.501 | **<0.001** |
| Basal respiration | Sp/Be-Be | Nutrient-poor | 0.546 | **<0.001** |
| Basal respiration | Sp-Be | Nutrient-poor | 0.642 | **0.002** |
| Microbial biomass | Do-Be | Nutrient-rich | 0.962 | 0.829 |
| Microbial biomass | Do/Be-Be | Nutrient-rich | 1.027 | 0.883 |
| Microbial biomass | Sp/Be-Be | Nutrient-rich | 0.825 | 0.286 |
| Microbial biomass | Sp-Be | Nutrient-rich | 0.946 | 0.756 |
| Microbial biomass | Do-Be | Nutrient-poor | 0.400 | **<0.001** |
| Microbial biomass | Do/Be-Be | Nutrient-poor | 0.428 | **<0.001** |
| Microbial biomass | Sp/Be-Be | Nutrient-poor | 0.413 | **<0.001** |
| Microbial biomass | Sp-Be | Nutrient-poor | 0.659 | **0.025** |
| Specific respiration | Do-Be | Nutrient-rich | 1.064 | 0.514 |
| Specific respiration | Do/Be-Be | Nutrient-rich | 0.985 | 0.869 |
| Specific respiration | Sp/Be-Be | Nutrient-rich | 1.151 | 0.144 |
| Specific respiration | Sp-Be | Nutrient-rich | 0.965 | 0.706 |
| Specific respiration | Do-Be | Nutrient-poor | 0.810 | **0.033** |
| Specific respiration | Do/Be-Be | Nutrient-poor | 1.020 | 0.833 |
| Specific respiration | Sp/Be-Be | Nutrient-poor | 0.890 | 0.223 |
| Specific respiration | Sp-Be | Nutrient-poor | 0.887 | 0.208 |
| Specific respiration | Do-Be | Nutrient-rich | 0.993 | 0.959 |
| cy/pre | Do-Be | Nutrient-rich | 1.080 | 0.592 |
| cy/pre | Do/Be-Be | Nutrient-rich | 1.062 | 0.674 |
| cy/pre | Sp/Be-Be | Nutrient-rich | 1.169 | 0.279 |
| cy/pre | Sp-Be | Nutrient-poor | 1.870 | **<0.001** |
| cy/pre | Do-Be | Nutrient-poor | 1.459 | **0.013** |
| cy/pre | Do/Be-Be | Nutrient-poor | 1.438 | **0.017** |
| cy/pre | Sp/Be-Be | Nutrient-poor | 1.395 | **0.027** |
| sat/mono | Do-Be | Nutrient-rich | 1.100 | 0.316 |
| sat/mono | Do/Be-Be | Nutrient-rich | 0.911 | 0.328 |
| sat/mono | Sp/Be-Be | Nutrient-rich | 1.058 | 0.555 |
| sat/mono | Sp-Be | Nutrient-rich | 1.070 | 0.479 |
| sat/mono | Do-Be | Nutrient-poor | 1.738 | **<0.001** |
| sat/mono | Do/Be-Be | Nutrient-poor | 1.372 | **0.002** |
| sat/mono | Sp/Be-Be | Nutrient-poor | 1.379 | **0.002** |
| sat/mono | Sp-Be | Nutrient-poor | 1.427 | **0.001** |
| Gram$^+$/Gram$^-$ | Do-Be | Nutrient-rich | 1.064 | 0.316 |
| Gram$^+$/Gram$^-$ | Do/Be-Be | Nutrient-rich | 0.989 | 0.860 |
| Gram$^+$/Gram$^-$ | Sp/Be-Be | Nutrient-rich | 1.038 | 0.542 |
| Gram$^+$/Gram$^-$ | Sp-Be | Nutrient-rich | 1.051 | 0.415 |
| Gram$^+$/Gram$^-$ | Do-Be | Nutrient-poor | 1.459 | **<0.001** |
| Gram$^+$/Gram$^-$ | Do/Be-Be | Nutrient-poor | 1.293 | **<0.001** |
| Gram$^+$/Gram$^-$ | Sp/Be-Be | Nutrient-poor | 1.227 | **0.002** |
| Gram$^+$/Gram$^-$ | Sp-Be | Nutrient-poor | 1.215 | **0.004** |

Table S2. Contrasts comparing the studied forest types (Douglas-fir [Do], Douglas-fir and European beech mixture [Do/Be], European beech [Be], Norway spruce and European beech mixture [Sp/Be] and Norway spruce [Sp]) at nutrient-poor sites (Poor) to nutrient-rich sites (Rich). Ratio was back-transformed from log response ratio. Significant p-values are given in bold ($p < 0.05$).

| Variable | Contrast | Forest type | Ratio | p-value |
|---|---|---|---|---|
| Basal respiration | Poor-Rich | Do | 0.431 | **0.004** |
| Basal respiration | Poor-Rich | Do/Be | 0.652 | 0.063 |
| Basal respiration | Poor-Rich | Be | 1.278 | 0.238 |
| Basal respiration | Poor-Rich | Sp/Be | 0.656 | 0.065 |
| Basal respiration | Poor-Rich | Sp | 0.972 | 0.884 |
| Microbial biomass | Poor-Rich | Do | 0.531 | **0.001** |
| Microbial biomass | Poor-Rich | Do/Be | 0.532 | **0.001** |
| Microbial biomass | Poor-Rich | Be | 1.276 | 0.179 |
| Microbial biomass | Poor-Rich | Sp/Be | 0.639 | **0.017** |
| Microbial biomass | Poor-Rich | Sp | 0.889 | 0.510 |
| Specific respiration | Poor-Rich | Do | 0.916 | 0.512 |
| Specific respiration | Poor-Rich | Do/Be | 1.246 | 0.133 |
| Specific respiration | Poor-Rich | Be | 1.203 | 0.195 |
| Specific respiration | Poor-Rich | Sp/Be | 0.930 | 0.585 |
| Specific respiration | Poor-Rich | Sp | 1.105 | 0.461 |
| cy/pre | Poor-Rich | Do | 1.817 | **0.017** |
| cy/pre | Poor-Rich | Do/Be | 1.303 | 0.198 |
| cy/pre | Poor-Rich | Be | 0.965 | 0.851 |
| cy/pre | Poor-Rich | Sp/Be | 1.306 | 0.194 |
| cy/pre | Poor-Rich | Sp | 1.151 | 0.471 |
| sat/mono | Poor-Rich | Do | 1.383 | **0.041** |
| sat/mono | Poor-Rich | Do/Be | 1.320 | 0.068 |
| sat/mono | Poor-Rich | Be | 0.876 | 0.331 |
| sat/mono | Poor-Rich | Sp/Be | 1.143 | 0.327 |
| sat/mono | Poor-Rich | Sp | 1.169 | 0.259 |
| Gram$^+$/Gram$^-$ | Poor-Rich | Do | 1.402 | **0.009** |
| Gram$^+$/Gram$^-$ | Poor-Rich | Do/Be | 1.336 | **0.017** |
| Gram$^+$/Gram$^-$ | Poor-Rich | Be | 1.022 | 0.816 |
| Gram$^+$/Gram$^-$ | Poor-Rich | Sp/Be | 1.208 | 0.078 |
| Gram$^+$/Gram$^-$ | Poor-Rich | Sp | 1.181 | 0.111 |
| Fungi | Poor-Rich | Do | 0.866 | 0.274 |
| Fungi | Poor-Rich | Do/Be | 1.085 | 0.531 |
| Fungi | Poor-Rich | Be | 1.303 | **0.049** |
| Fungi | Poor-Rich | Sp/Be | 1.265 | 0.078 |
| Fungi | Poor-Rich | Sp | 1.345 | **0.029** |
| Bacteria | Poor-Rich | Do | 0.972 | 0.449 |
| Bacteria | Poor-Rich | Do/Be | 1.012 | 0.754 |
| Bacteria | Poor-Rich | Be | 0.961 | 0.293 |
| Bacteria | Poor-Rich | Sp/Be | 0.961 | 0.297 |
| Bacteria | Poor-Rich | Sp | 0.925 | **0.046** |
| Fun/Bac | Poor-Rich | Do | 0.868 | 0.404 |
| Fun/Bac | Poor-Rich | Do/Be | 1.067 | 0.697 |
| Fun/Bac | Poor-Rich | Be | 1.292 | 0.156 |
| Fun/Bac | Poor-Rich | Sp/Be | 1.271 | 0.180 |
| Fun/Bac | Poor-Rich | Sp | 1.449 | 0.058 |

Table S3. Forward selection in redundancy analyses (RDA). Statistically significant covariates were determined using stepwise forward selection after Monte Carlo permutation test. P-values < 0.05 and significant accumulated $R^2$ values are given in bold.

| Depth | Based on P-values (“ordistep”) | | | Based on adjusted $R^2$ (“ordiR2step”) | |
|---|---|---|---|---|---|
| | Tested term | F | P (>F) | Forward selected model | Adjusted $R^2$ |
| Litter | pH | 4.45 | **0.005** | = pH | **0.08** |
| | C/N | 1.65 | 0.090 | = pH + C/N | 0.10 |
| | Water% | 1.81 | 0.105 | = pH + Water% | 0.10 |
| | Litter mass | 1.35 | 0.215 | = pH + Litter mass | 0.09 |
| | C% | 1.06 | 0.395 | = pH + C% | 0.08 |
| | | | | All terms | 0.13 |
| 0–5 cm | C/N | 11.76 | **0.005** | = C/N | **0.22** |
| | pH | 5.61 | **0.005** | = C/N + pH | **0.30** |
| | Water% | 4.79 | **0.005** | = C/N + pH + Water% | **0.37** |
| | Litter mass | 0.80 | 0.610 | = C/N + pH + Water% + Litter mass | 0.36 |
| | C% | 1.20 | 0.300 | = C/N + pH + Water% + C% | 0.37 |
| | | | | All terms | 0.37 |
| 5–10 cm | C/N | 7.01 | **0.005** | = C/N | **0.13** |
| | C% | 1.82 | 0.075 | = C/N + C% | 0.15 |
| | Water% | 1.19 | 0.260 | = C/N + Water% | 0.14 |
| | pH | 0.81 | 0.580 | = C/N + pH | 0.13 |
| | Litter mass | 0.99 | 0.435 | = C/N + Litter mass | 0.13 |
| | | | | All terms | 0.15 |
